# TAL effectors with avirulence activity in African strains of *Xanthomonas oryzae* pv. *oryzae*

**DOI:** 10.1101/2021.10.21.465293

**Authors:** Marlène Lachaux, Emilie Thomas, Adam J. Bogdanove, Boris Szurek, Mathilde Hutin

**Affiliations:** PHIM, IRD, CIRAD, INRAe, Institut Agro, Univ. Montpellier, Montpellier, France; Plant Pathology and Plant Microbe Biology Section, School of Integrative Plant Science, Cornell University, Ithaca, NY, 14853 United States of America

**Keywords:** Rice, bacterial leaf blight, truncTALE/iTALE, germplasm, executor gene, *Xa1*-like resistance

## Abstract

**Background:** *Xanthomonas oryzae* pv. *oryzae* causes bacterial leaf blight, a devastating disease of rice. Among the type-3 effectors secreted by *Xanthomonas oryzae* pv. *oryzae* to support pathogen virulence, the Transcription Activator-Like Effector (TALE) family plays a critical role. Some TALEs are major virulence factors that activate susceptibility (*S*) genes, overexpression of which contributes to disease development. Host incompatibility can result from TALE-induced expression of so-called executor (*E*) genes leading to a strong and rapid resistance response that blocks disease development. In that context, the TALE functions as an avirulence (Avr) factor. To date no such avirulence factors have been identified in African strains of *Xanthomonas oryzae* pv. *oryzae*.

**Results:** With respect to the importance of TALEs in the Rice-*Xoo* pathosystem, we aimed at identifying those that may act as Avr factor within African *Xoo*. We screened 86 rice accessions, and identified 12 that were resistant to two African strains while being susceptible to a well-studied Asian strain. In a gain of function approach based on the introduction of each of the nine *tal* genes of the avirulent African strain MAI1 into the virulent Asian strain PXO99^A^, four were found to trigger resistance on specific rice accessions. Loss-of-function mutational analysis further demonstrated the *avr* activity of two of them, *talD* and *talI*, on the rice varieties IR64 and CT13432 respectively. Further analysis of TalI demonstrated the requirement of its activation domain for triggering resistance in CT13432. Resistance in 9 of the 12 rice accessions that were resistant against African *Xoo* specifically, including CT13432, could be suppressed or largely suppressed by trans-expression of the truncTALE *tal2h*, similarly to resistance conferred by the *Xa1* gene which recognizes TALEs generally independently of their activation domain.

**Conclusion:** We identified and characterized TalD and TalI as two African *Xoo* TALEs with avirulence activity on IR64 and CT13432 respectively. Resistance of CT13432 against African *Xoo* results from the combination of two mechanisms, one relying on the TalI-mediated induction of an unknown executor gene and the other on an *Xa1*-like gene or allele.

## Introduction

Cultivated plants constantly face multiple abiotic and biotic stresses, the latter of which are estimated to cause from 17 to 30% of global yield losses on five of the most important crops including rice (Savary et al. 2019). Rice (*Oryza sativa L*.) is one of the most widely cultivated crops around the world and a staple food for much of the developing world (Ainsworth 2008). A major threat to rice production in Asia and Africa is bacterial leaf blight (BLB) caused by the bacterial phytopathogen *Xanthomonas oryzae* pv. *oryzae* (*Xoo*). BLB may indeed cause up to 50% of yield loss depending on rice variety, growth stage of infection, geographic location and environmental conditions (Liu et al. 2014). *Xoo* enters leaves through hydathodes or wounds. Bacteria then multiply in the intercellular spaces of the underlying epitheme prior to reaching the xylem vessels and propagating into the plant. BLB symptoms are water-soaked lesions that spread following the bacteria’s progression down the leaf and become chlorotic and then necrotic (Niño-Liu et al. 2006).

As do many pathogenic gram-negative bacteria, *Xoo* uses a type-3 secretion system (T3SS) to secrete into the host cytoplasm a cocktail of type-3 effectors (T3E) that can be classified as transcription activator-like (TAL) effectors and non-TAL effectors. While the latter include a diverse array of effector families with various molecular activities, members of the TAL effector (TALE) family function as eukaryotic transcription factors that bind in a sequence-specific manner to the promoters of target genes in the host cells. Repeat-variable diresidues (RVDs) located at positions 12^th^ and 13^th^ of each of several contiguous repeats in a central domain of TALEs determine the DNA sequence binding specificity of the protein, one repeat to one nucleotide (Boch et al. 2009; Moscou and Bogdanove 2009). The target sequence, unique to each TALE, is called the effector binding element (EBE). Some *Xoo* TALEs are major virulence factors, targeting susceptibility (*S*) genes. TALE driven upregulation of these genes contributes to disease development. *S* genes characterized to date for BLB mainly encode clade-3 *SWEET* sugar uniporters that may increase the abundance of apoplastic sugar to the benefit of the pathogen, or transcription factors that regulate so far unknown secondary targets promoting host susceptibility (Garcia-Ruiz et al. 2021).

Forty-six genes, several dominant and some recessive, individually govern rice resistance against *Xoo*. Twelve have been cloned and nine are TALE-dependent, reflecting the crucial role of TALEs in the interaction (Jiang et al. 2020). Recessive resistance often involves mutations within the EBE of an *S* gene to prevent the TALE-DNA interaction and consequent *S* gene induction. This is well illustrated by the non-TALE-inducible *xa13, xa25* and *xa41* loss-of-susceptibility alleles of the major *S* genes *OsSWEET11, OsSWEET13*, and *OsSWEET14*, respectively (Chu et al. 2006; Liu et al. 2011; Hutin et al. 2015). Dominant resistance is often triggered by TALE-mediated induction of so-called executor (*E*) genes, expression of which leads to rapid plant cell death that blocks disease development (Boch et al. 2014). Four *E* genes, namely *Xa7, Xa10, Xa23, Xa27*, and their respective matching *tal* genes, *avrXa7, avrXa10, avrXa23, avrXa27*, have been cloned and characterized (Hopkins et al. 1992; Gu et al. 2005; Tian et al. 2014; Wang et al. 2014, 2015; Chen et al. 2021; Luo et al. 2021). These *tal* genes are only present in Asian strains of *Xoo* and no *E* genes induced by African *Xoo* TALEs have been identified to date. *E* genes so far code for small proteins with transmembrane domains, and the molecular mechanisms underlying their function are still unclear far from being understood (Zhang et al. 2015; Chen et al. 2021). Another type of dominant resistance is conferred by receptor-like kinases (RLK), such as *Xa3/Xa26* (Sun et al. 2004; Xiang et al. 2006) and *Xa21* (Song et al. 1995), but also *Xa4* which encodes a cell wall-associated kinase (Hu et al. 2017). Finally, the last category are genes encoding nucleotide-binding domain leucine-rich repeat containing receptors (NLR) such as *Xa1* and its apparent alleles *Xo1, Xa2, Xa14, Xa31(t)* and *Xa45(t)* (Ji et al. 2020; Read et al. 2020; Zhang et al. 2020). We consider the latter genes “apparent” alleles because *Xo1* resides in a cluster of NLR genes, the number of which varies among rice genotypes, making orthology uncertain. Xa1 and Xo1 were shown to mediate resistance in response to TALEs generally, independently of their specific RVD sequence, and with no requirement for the transcriptional activation domain (Ji et al. 2016; Triplett et al. 2016; Read et al. 2020). A variant class of TALEs called interfering (iTALE) or truncated (truncTALE) TALEs suppress *Xa1*/*Xo1*-mediated resistance, and the truncTALE Tal2h has been demonstrated to interact with Xo1 (Ji et al. 2016; Read et al. 2016, 2020). Whether this interaction is direct, and whether Xo1 interacts with TALEs to mediate resistance remains to be elucidated. The analysis of functional apparent alleles of *Xa1* and *Xo1* highlight the systematic absence of an intervening motif present in non-functional alleles, and differences in the number (4 to 7) of central tandem repeats that might explain differences in their activity (Zhang et al. 2020). Interestingly, most Asian strains of *Xoo* harbor iTALEs/truncTALEs while African strains do not, explaining why African strains are widely controlled by *Xa1, Xo1* or functional homologs (Ji et al. 2016; Read et al. 2016, 2020).

Previous studies demonstrated that African *Xoo* are genetically distant from Asian *Xoo* and closer to *Xanthomonas oryzae* pv. *oryzicola* (*Xoc*) (Poulin et al. 2015), which causes bacterial leaf streak. A characteristic feature of African *Xoo* is their small TALome (set of *tal* genes) consisting of 8-9 genes, relative to Asian *Xoo* which carry up to 19 *tal* genes. Moreover, no TALE is conserved between Asian and African strains (Lang et al. 2019). Comparative analysis of the TALome of several African strains revealed six groups of polymorphic TALEs based on their RVD sequences, including TalA, TalB, TalD, TalF, TalH, and TalI (Doucouré et al. 2018; Tran et al. 2018). African *Xoo* are also distinguished by a reduced number of races as compared to Asian *Xoo* (Gonzalez et al. 2007; Tekete et al. 2020). Race profiling on near isogenic lines (NILs) reported the potential of *Xa4, xa5* and *Xa7* or co-segregating genes to control a few African *Xoo* from Burkina Faso, Niger, and Cameroon (Gonzalez et al. 2007), but the resistance spectrum of these genes has yet to be evaluated on a larger set of strains. Furthermore, *Xa1* comes up as one of the most promising *R* genes in terms of resistance spectrum for the Malian *Xoo* population (Tekete et al. 2020). Other studies to identify resistance against African *Xoo* evaluated 107 accessions of *O. glaberrima*, the cultivated rice species domesticated in Africa, as well as improved varieties including NERICA (NEw RICe for Africa) (Djedatin et al. 2011; Wonni et al. 2016). NERICA varieties are often the result of the inter-specific crosses between *O. glaberrima*, which represents important germplasm for resistance to local biotic and abiotic stresses, and high-yielding Asian *O. sativa*. These studies identified 25 accessions of *O. glaberrima* that are resistant to one or more African strains of *Xoo*, and five Burkinabe elite rice varieties (Djedatin et al. 2011; Wonni et al. 2016). Genes or quantitative trait loci accounting for resistance in these varieties remain to be explored.

In this study, toward providing breeders with new resistance genes against African *Xoo*, we screened 86 rice accessions including 16 accessions tested previously (Djedatin et al. 2011; Wonni et al. 2016) for resistance to two reference African *Xoo* strains MAI1 and BAI3, respectively originating from Mali and Burkina Faso. We included the Asian strain PXO99^A^ for comparison. For select accessions, we probed with individual TALEs from the African strains expressed in PXO99^A^ to identify potential *E* or other TALE-dependent resistance genes. We report on the identification of 12 accessions showing resistance to both African strains, nine of which involving an *Xa1*-like immunity, and unveil two TALEs with avirulence activity in African *Xoo*. Interestingly, our approach unmasked the occurrence of two overlapping TALE-mediated sources of resistance in the rice variety CT13432, one involving Xa1-like activity and the other a so far unknown TalI-dependent *E* gene.

## Results

### Germplasm screening for TALE-dependent resistance against African *Xoo* uncovers three resistant rice varieties

To search for African *Xoo tales* with *avr* activity, we established a gain-of-function approach consisting in the trans-expression of these *tal* genes in a virulent recipient strain of *Xoo*. We first screened a germplasm of 86 accessions of rice and selected those that were susceptible to the Asian *Xoo* strain PXO99^A^ and resistant to the reference African *Xoo* strains MAI1 and BAI3. Twelve accessions exhibited that phenotype, including two *O. glaberrima*, three *O. sativa* (two *indica* and one *japonica*), and seven elite varieties that are popular in West-Africa (Table S1). To investigate whether the resistance of these 12 accessions to African *Xoo* is triggered by TALEs, each of the nine *tal* genes of the *Xoo* strain MAI1 was introduced into the virulent strain PXO99^A^. Each PXO99^A^ transformant carrying an *Xoo* MAI1 *tal* gene was inoculated to the 12 accessions and to the rice variety Azucena, which was used as susceptible check. No significant difference in lesion lengths was observed upon leaf-clip inoculation of Azucena leaves with the different transformants 15 days after inoculation. In contrast, the varieties CT13432 and FKR47N exhibited resistance when inoculated with PXO99^A^ transformants carrying *talI* and *talF*, respectively. In addition, PXO99^A^ strains with *talD* or *talH* both elicited resistance when inoculated to the *O. sativa* ssp. *indica* variety IR64. Overall, four TALEs with avirulence activity and three rice accessions with TAL-dependent resistance were pinpointed through this gain-of-function strategy (Table 1).

**Table 1.** Four African TALEs mediate resistance when expressed in PXO99^A^ on resistant varieties.

| Name | Species | MAI1<sup>°</sup> | BAI3<sup>°</sup> | PXO99<sup>Ao</sup> | *Candidate avirulence TALEs |
| --- | --- | --- | --- | --- | --- |
| Og_107 | *O. glaberrima* | R | MR | S | - |
| Og_162 | *O. glaberrima* | R | MR | S | - |
| WAB56-50 | *O. sativa* spp. *japonica* | R | R | S | - |
| WAB181-18 | *O. sativa* spp. *japonica* | R | R | S | - |
| FKR19 | *O. sativa* spp. *japonica* | R | R | S | - |
| FKR43 | *O. sativa* spp. *japonica* | R | R | S | - |
| IR64 | *O. sativa* spp. *indica* | R | MR | S | TalD, TalH |
| CT13432 | *O. sativa* spp. *japonica* | R | R | S | TalI |
| Gigante | *O. sativa* | R | R | S | - |
| FKR45N | *O. sativa* spp. *japonica* / *O. glaberrima* | R | R | S | - |
| FKR47N | *O. sativa* spp. *japonica* / *O. glaberrima* | R | R | S | TalF |
| FKR49N | *O. sativa* spp. *japonica* / *O. glaberrima* | R | R | S | - |

<sup>°</sup>Resistance or susceptibility of rice to *Xoo* is expressed as a result of lesion length measurements 15 days after inoculation. Resistant (R); moderately resistant (MR); moderately susceptible (MS); susceptible (S).
\*Rice accessions were inoculated with PXO99<sup>A</sup> carrying each of the nine *tales* of MAI1. (-) means no candidate avirulence TALE identified.

### *tal* genes mutagenesis confirms *talD* avirulence activity in IR64

To confirm that *talD, talH, talI* and *talF* act as avirulence genes in strain MAI1, we attempted to generate a library of MAI1 *tal* gene mutants by transformation of the suicide plasmid pSM7 as reported previously (Cernadas et al. 2014; Tran et al. 2018). Because MAI1 turned out to be poorly amenable to genetic transformation, we focused on the *Xoo* strain BAI3, which has a similar TALome (Tran et al. 2018). We obtained at least one mutant strain with a single insertion for each *tal* gene, except for *talH* (Fig. S1). Alongside wild-type (WT) BAI3, mutant strains BAI3Δ*talI*, BAI3Δ*talF* and BAI3Δ*talD* were inoculated to CT13432, FKR47N and IR64, respectively, and to the Azucena susceptible control. Both leaf-clip inoculation and leaf-infiltration of the three BAI3Δ*tal* mutants produced WT-like symptoms on Azucena, indicating that virulence of these mutant strains was not affected (Fig. 1; Fig. S2). In contrast, an increase of lesion lengths was observed upon clip-inoculation of leaves of IR64 with BAI3Δ*talD*. Avirulence was fully restored when a plasmid-borne copy of *talD* was introduced into BAI3Δ*talD* (Fig. 1A), demonstrating the avirulence activity of *talD* in IR64. In contrast, no loss of resistance was observed upon leaf-infiltration or leaf-clipping of CT13432 and FKR47N with *talI* or *talF* mutant strains, respectively (Fig. S2, Fig. 1B), leading to the hypothesis that one or more other *avr* activities, corresponding to one or more other *R* genes, may be masking those of *talI* and *talF*.

**Fig. 1.**
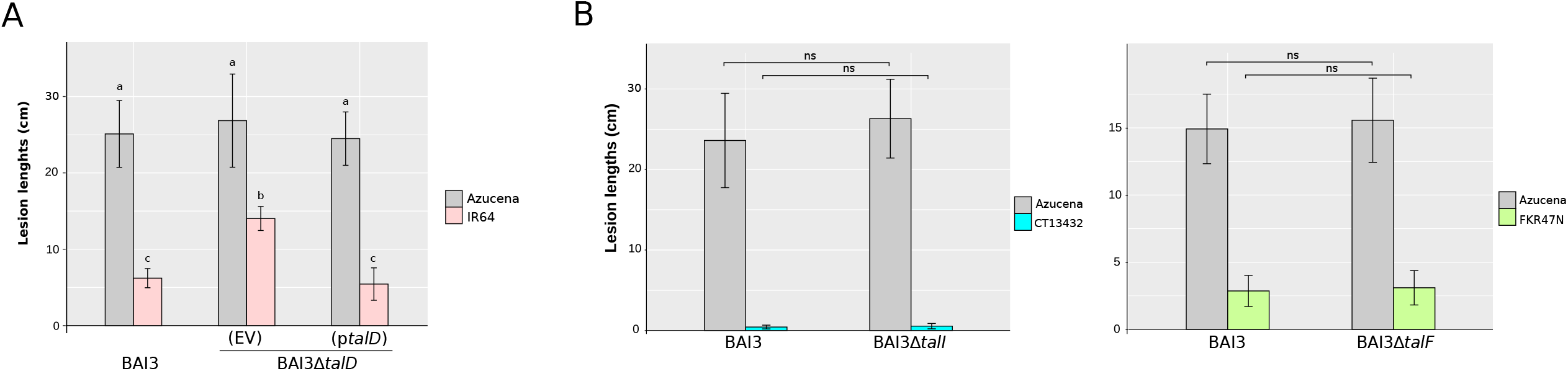
BAI3Δ*talD* loses avirulence on IR64 but BAI3Δ*talI* and BAI3Δ*talF* do not on CT13432 and FKR47N. (A) Leaves of IR64 and Azucena were challenged with *Xoo* strains BAI3, and BAI3Δ*talD* carrying an empty vector (EV) or a plasmid with *talD* (p*talD)*. To assess significance, the parametrical test ANOVA (α = 0.05) was carried out using the R package. Values labeled with the same lower-case letter are not significantly different. (B) Leaves of rice varieties CT13432, and FKR47N were clip-inoculated with the wild-type African *Xoo* strain BAI3 and with mutant strains BAI3Δ*talI* and BAI3Δ*talF*, respectively. Azucena was included for all strains as a susceptible check. Lesion lengths were measured at 15 dpi. Data are the mean of eight measurements. Error bars represent ± SD. Significant differences (p < 0.001) were determined using an unpaired two-sample t-test, bars with « ns » are not statistically different. Experiments were repeated three time with similar results.

### CT13432 and FKR47N exhibit *Xa1*-like resistance

When leaves of CT13432 and FKR47N were infiltrated with BAI3, an early and strong hypersensitive response (HR) apparent as rapidly developing necrosis could be observed at the site of inoculation (Fig. S2). *Xa1*/*Xo1* being able to confer resistance against African strains of *Xoo* and *Xoc* specifically (Ji et al. 2016; Triplett et al. 2016), and among the *Xa* genes that trigger an early and strong HR-like phenotype upon *Xoo* leaf-infiltration, we hypothesized that some *Xa1/Xo1*-like mechanism may be at play in the BAI3-CT13432/FKR47N interactions, redundant to the *talI* and *talF-*specific resistances observed in the gain-of-function experiments. To test this hypothesis we infiltrated the Asian *Xoc* strain BLS256 which carries the truncTALE *tal2h*, and the derivative mutant strain BLS256Δ*tal2h*, into leaves of CT13432, FKR47N and the susceptible control Azucena (Fig. 2). As expected, all strains promoted water-soaking symptoms upon infiltration of the susceptible variety Azucena. Typical BLS symptoms were observed when leaves of CT13432 and FKR47N were infiltrated with *Xoc* strain BLS256, while a strong resistance phenotype appeared upon infiltration of the truncTALE derivative mutant BLS256Δ*tal2h*, indicating that CT13432 and FKR47N have *Xa1*/*Xo1*-like activity. To further confirm that the resistance of these varieties against *Xoo* strains BAI3 and MAI1 is conferred in part by an *Xa1*-like gene, both strains were transformed with *tal2h* and pathogenicity assays were performed. As expected, expression of the truncTALE in BAI3 and MAI1 resulted in a complete loss of HR on CT13432 and FKR47N, which was not the case when the strains carried an empty vector (Fig. 2). Altogether, our results suggest that African *Xoo* resistance in CT13432 and FKR47N is mediated by *Xa1* or other alleles, in addition to as yet unidentified resistance genes corresponding to *talI* and *talF*.

**Fig. 2.**
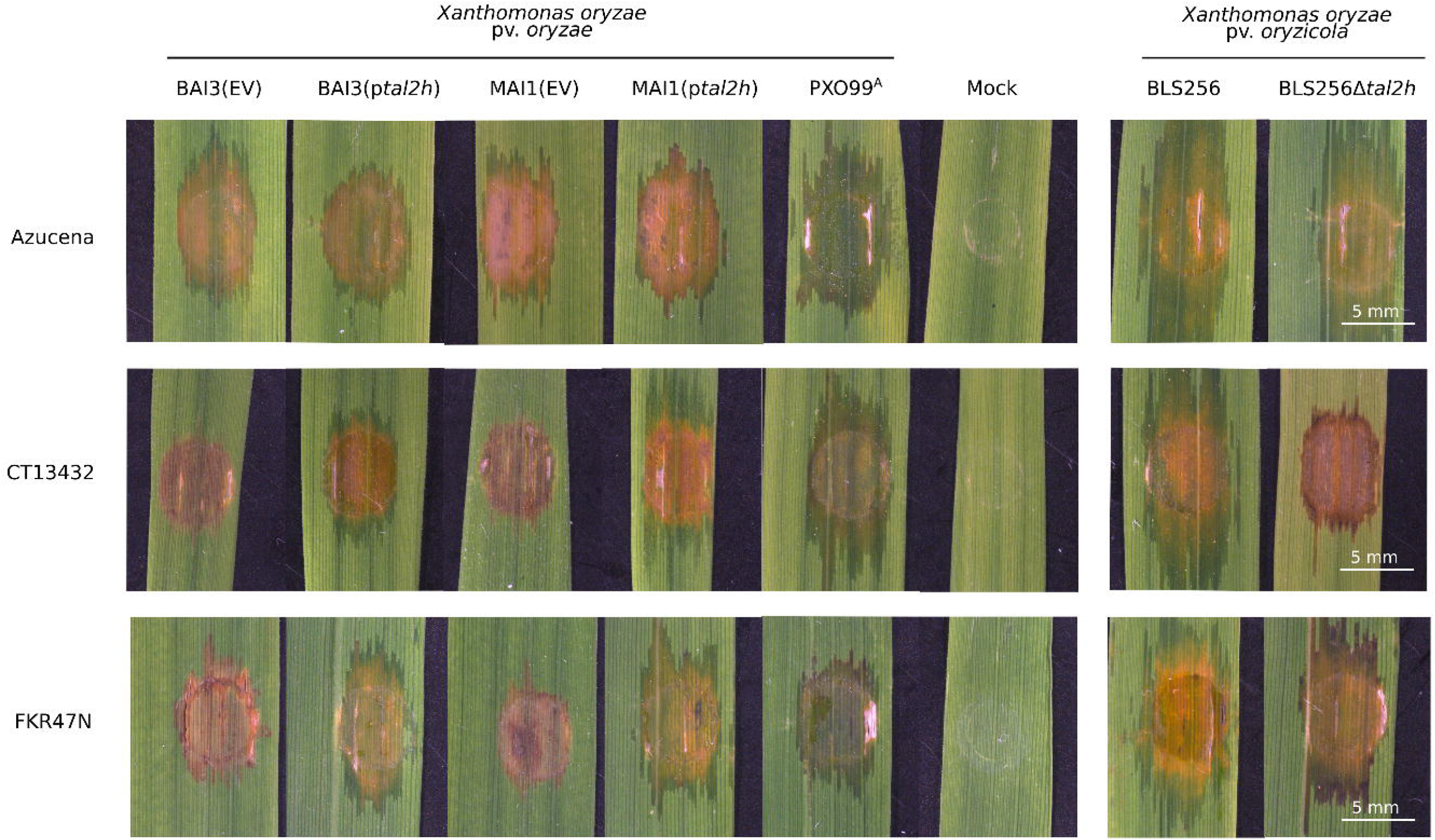
The resistance of CT13432 and FKR47N against African *Xoo* is largely suppressed by a truncTALE. Leaves of CT13432 and FKR47N rice varieties were inoculated with *Xoo* strains BAI3 and MAI1 carrying an empty vector (EV) or the truncTALE *tal2h* as well as *Xanthomonas oryzae* pv. *orizycola* (*Xoc*) strain BLS256 which is naturally carrying *tal2h* and the mutant strain BLS256Δ*tal2h*. The susceptible rice variety Azucena and the Asian *Xoo* strain PXO99^A^ were used as controls. Leaves were photographed at 5 dpi.

To test whether any of the ten remaining resistant accessions involve similar mechanisms, 3-week-old plants were inoculated by infiltration and checked for appearance of the HR. As expected, Carolina Gold Select and IRBB1, carrying respectively *Xo1* and *Xa1*, displayed water-soaking lesions when infiltrated with the Asian *Xoc* strain BLS256, and HR when infiltrated with the African *Xoo* strain BAI3 and the truncTALE derivative mutant BLS256Δ*tal2h* (Fig. S3). Surprisingly, seven out of the eight *O. sativa* spp. accessions also exhibited a typical *Xa1*-like HR to BAI3 and BLS256Δ*tal2h* while being susceptible to BLS256 (and to PXO99^A^, which also carries iTALEs). The two *O. glaberrima* accessions exhibited no Xa1-like resistance, developing water-soaking in response to BAI3 or BLS256Δ*tal2h* following leaf infiltration (Fig. S3, Table S2). Overall, these observations show that 9 of the 12 resistant varieties including CT13432 and FKR47N resist African *Xoo* through *Xa1/Xo1*-like mechanisms.

### Presence of *Xa1* allelic *R* genes in CT13432 and FKR47N rice varieties

To test the prediction that an *Xa1* allele was present in CT13432 and FKR47N, we PCR-amplified a 202 bp fragment spanning the junction of the first repeat and its up-stream region (Ji et al. 2020); Table S3), allowing the discrimination of resistant and susceptible alleles (susceptible alleles yield no product). Analysis was performed also on the *Xa1*-carrying near-isogenic line IRBB1 and the *Xo1*-carrying Carolina Gold Select rice variety as positive controls, and on Azucena, Nipponbare and IR24 as negative controls. Both CT13432 and FKR47N yielded a product that co-migrated with those of the positive controls (Fig. 3A). No amplification was evident from the negative control varieties. We next tried to determine the number of leucine-rich repeats (LRRs) encoded by the CT13432 and FKR47N alleles by amplifying the LRR domain. As expected, IRBB1 and Carolina Gold yielded products of sizes consistent with the six and five LRRs of Xa1 and Xo1, respectively. CT13432 and FKR47N produced amplicons of sizes consistent with the presence of seven and five LRRs, respectively (Fig. 3B). Altogether, these results provide strong evidence that CT13432 and FKR47N varieties each carry a different allele of the *Xa1 R* gene that is active against African *Xoo* strains MAI1 and BAI3.

**Fig. 3.**
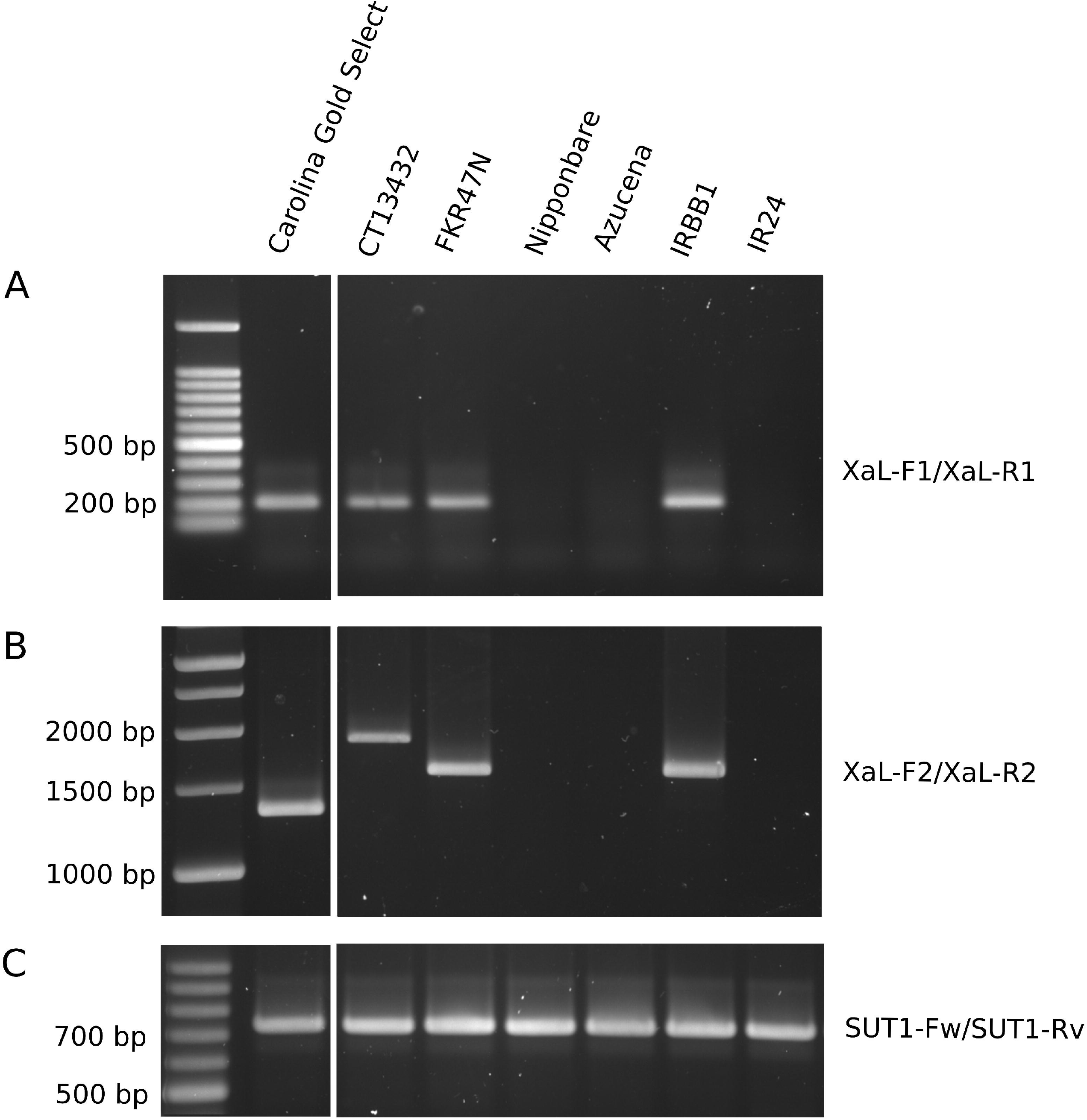
The rice varieties CT13432 and FKR47N carry distinct apparent alleles of *Xa1*. The genomic DNA of rice varieties Carolina Gold Select, CT13432, FKR47N, Nipponbare, Azucena, IRBB1 and IR24 were used as a template for PCR amplification of (A) the junction of the first repeat of 279 base pairs of *Xa1* and its upstream region, (B) the whole repeat region of *Xa1* and alleles, and (C) the housekeeping gene *OsSUT1* used as a control. All the samples were run on the same gel.

### Suppression of the *Xa1*-like resistance in CT13432 unmasks an underlying TalI activation domain-dependent resistance

We hypothesized that the *Xa1-*like resistance in CT13432 and FKR47N explains the failure of the *talI* and *talF* knockouts in BAI3 to abolish avirulence of the mutants, respectively, on these varieties (Fig. 1A). To test this hypothesis directly, we first introduced a plasmid-borne copy of *tal2h* into BAI3 and inoculated to CT13432 and the susceptible variety Azucena. The presence of *tal2h* indeed suppressed the *Xa1*-like resistance to BAI3 in CT13432, resulting in moderate disease symptoms development (Fig. 4A), and higher bacterial titers *in planta* (Fig. 4B) as compared with a BAI3 empty vector control. We next tested whether the combination of *tal2h*-mediated *Xa1*-like suppression and *talI* inactivation would lead to even higher virulence on CT13432 by introducing *tal2h* (or the empty vector) into BAI3Δ*talI*(p*tal2h*). No significant differences in lesion lengths or bacterial populations were observed between BAI3(p*tal2h*) and BAI3Δ*talI*(p*tal2h*)(EV). However, expression of *talI*_*BAI3*_ in trans in BAI3Δ*talI*(p*tal2h*) dramatically reduced lesion lengths and bacterial titer *in planta*, relative to the empty vector in BAI3Δ*talI*(p*tal2h*). These results demonstrate that *talI*_*BAI3*_ *avr* activity in CT13432 can be detected when *Xa1* is inactivated. We next evaluated if restoration of avirulence could also be obtained upon expression of *talI*_*MAI1*_ which is polymorphic at RVDs 4 and 9 (Tran et al. 2018). As with *talI*_*BAI3*_, expression of *talI*_*MAI1*_ lead to reduced lesion lengths and *in planta* bacterial amounts, indicating that both *talI* variants confer avirulence on CT13432 (Fig. 4). We next evaluated the requirement of the activation domain (AD) in the *talI*-specific resistance. BAI3Δ*talI*(p*tal2h*) transformed with the deletion derivative construct p*talI*_*MAI1*_ΔAD failed to trigger resistance in CT13432 while it remained fully virulent on the susceptible variety Azucena (Fig. 4A). Western-blot analysis verified that TalI_MAI1_ and TalI_MAI1_ΔAD accumulate at comparable levels (Fig. S4). Altogether our results show that the TalI specifically elicits a form of resistance distinct from the Xa1-like resistance in CT13432 and that this elicitation relies on the activation domain of TalI, suggesting the presence of a TalI-activated *E* gene. Further analysis will be necessary to elucidate whether the same is true for *talF-*mediated resistance in FKR47N.

**Fig. 4.**
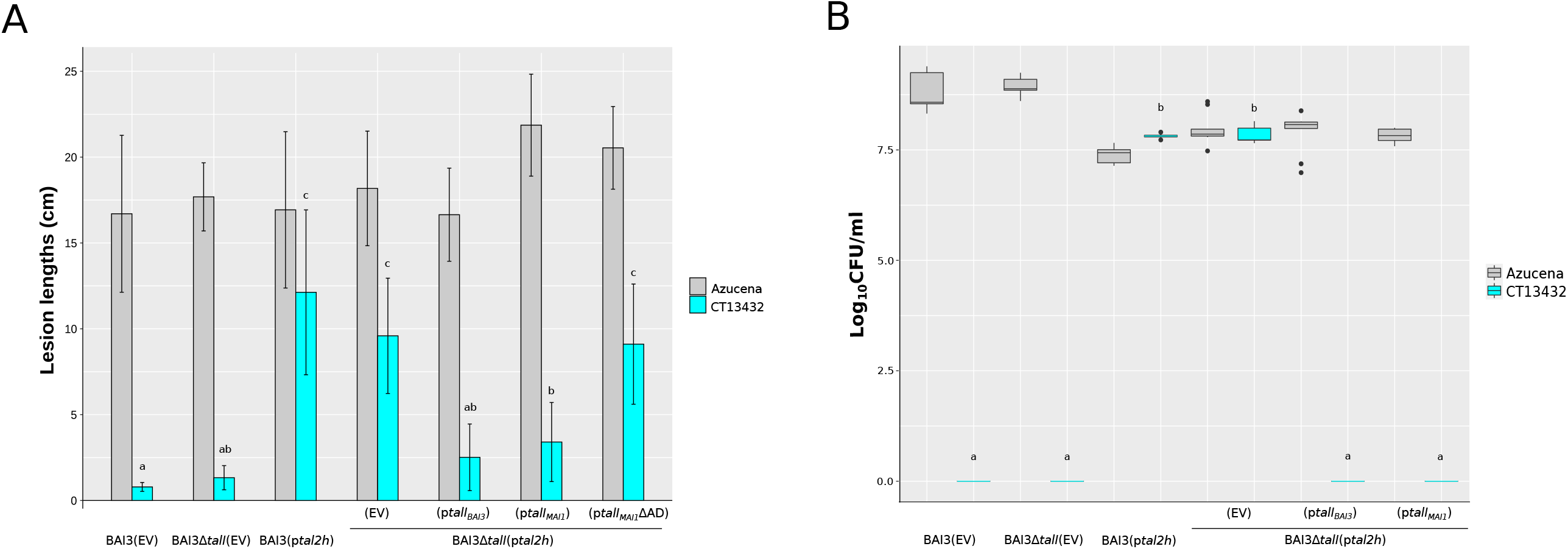
Suppression of Xa1-like resistance unmasks *talI* avirulence activity in the rice variety CT13432. (A) Leaves of CT13432 and Azucena plants were clip-inoculated with the *Xoo* strain BAI3 carrying the pKEB31 empty vector (EV), the BAI3Δ*talI* mutant carrying the pKEB31 empty vector (EV), the BAI3 strain expressing the *tal2h* truncTALE, and BAI3Δ*talI* expressing *tal2h* and the pSKX1 empty vector (EV), or pSKX1 containing *talI*_*BAI3*_, *talI*_*MAI1*_ or *talI*_*MAI1*_ΔAD. Lesion lengths were measured at 15 dpi. Data are the mean of eight measurements. (B) Bacterial counts were taken 7 days post-inoculation of rice leaves of CT13432 and Azucena inoculated with the same panel of *Xoo* strains used in (A). Statistical tests in both experiments were carried out in R using the nonparametric Kruskal-Wallis and Dunn’s tests (α = 0.05). Values labeled with the same lower-case letter are not significantly different.

## Discussion

First identified in Japan in 1884, BLB has been long recognized as a major threat in many Asian countries, and therefore for decades at the heart of many research and resistance breeding programs (Liu et al. 2014). To date, a handful of avirulence genes have been cloned from Asian *Xoo* strains, all of which encode TAL effectors including *avrXa7, avrXa10, avrXa23*, and *avrXa27* (Hopkins et al. 1992; Gu et al. 2005; Wang et al. 2014). In contrast, no avirulence gene has been identified in African *Xoo* strains, the first of which were isolated in the 1980s. However, as many as nine races of African strains, not found among Asian strains, have been reported so far (Gonzalez et al. 2007; Tekete et al. 2020), indicating that some effector genes of African *Xoo* strains act as avirulence factors in gene-for-gene interactions with some rice *R* genes. Comparison between African and Asian *Xoo* TALomes shows that no *tal gene* is conserved between these two *Xoo* lineages (Lang et al. 2019). Strains of both lineages induce the *S* genes *OsSWEET14* and *OsTFX1* through unrelated TALEs, highlighting cases of inter-lineage evolutionary convergence of virulence functions (Streubel et al. 2013; Tran et al. 2018). In contrast, none of the rice *E* genes induced by Asian *Xoo* is predicted to be targeted by any African *Xoo* TALE. Here, toward identifying sources of resistance effective against African *Xoo* strains, we assessed the putative *avr* activity of 12 previously cloned individual *tal* genes representing the TALome of two African *Xoo* strains (Tran et al. 2018), using a large set of rice accessions.

To accomplish this goal, we first took a gain-of-function approach using the virulent Asian *Xoo* strain PXO99^A^ as a recipient to express cloned African *tal* genes. In a similar approach, we previously identified major virulence TALEs of African *Xoo* strains using the US strain *Xo* X11-5A as recipient (Tran et al. 2018). X11-5A lacks *tal* genes and is weakly virulent, allowing for gain-of-virulence screening (Ryba-White et al. 1995; Triplett et al. 2011). Nonetheless in a search for gain-of-avirulence, the use of a more virulent strain is required. By selecting the Asian strain PXO99^A^ as recipient, we also minimized the risk of TALE functional redundancy. This strategy led to the identification of *talI* and *talF* respectively causing resistance in rice varieties CT13432 and FKR47N, as well as *talD* and *talH*, each triggering resistance on IR64.

In a complementary approach, a loss-of-function analysis was carried out by mutagenizing individual *tal* genes in the African *Xoo* strain BAI3. This analysis validated *talD* and *talI* as avirulence genes; a mutant for *talH* was not obtained. Because IR64 has been sequenced and recombinant populations are available, (Fragoso et al. 2017) prospects for future identification of the gene(s) underlying the TalD-elicited resistance are good. Mutagenesis of *talI* or *talF* failed to cause a measurable loss of avirulence in the respective rice varieties CT13432 and FKR47N. However, *talI* avirulence activity was evident upon trans-expression of the truncTALE *tal2h* in BAI3Δ*talI*, revealing that a resistance like that mediated by Xa1, in response to TALEs generally and suppressed by truncTALEs, is present in CT13432 and masking the TalI-specific resistance. By further taking advantage of this combined gain- and loss-of-function approach, we also determined that the TalI-specific resistance depends on the activation domain of the effector and thus likely involves an *E* gene.

While four TALE groups including TalG, TalE, TalD and TalC are strictly conserved between *Xoo* strains MAI1 and BAI3, two to six RVD variations were reported in the five other groups, modifying to some extent their DNA binding sequence specificities (Tran et al. 2018). Notably, three of the four candidate or validated avirulence TALEs identified in this study, TalI, TalF, and TalH all present RVD polymorphisms across Malian and Burkinabe strains, which could be the result of some ongoing selection pressure imposed by corresponding *E* or other resistance genes (Doucouré et al. 2018; Schandry et al. 2018). Surprisingly, the TalD group is much more conserved, highlighted by an absolute RVD conservation between BAI3, MAI1 and the Cameroonian strain CFBP1947 (Tran et al. 2018), and only one polymorphic RVD among nine Malian strains investigated (Doucouré et al. 2018). According to the EBE prediction tools Target Finder and Talvez (Doyle et al. 2012; Pérez-Quintero et al. 2013), 90% of the first 20 predicted targets of the two versions of TalD are conserved. Such a conservation among strains hints to an important role in virulence for TalD, but there are no data supporting this hypothesis so far. Concerning the TalI group, BAI3 and MAI1 alleles differ at two RVDs and share about 32% of their predicted targets, querying the Nipponbare genome. Since both variants equally elicit resistance in the rice variety CT13432, their polymorphism may confer an advantage in future identification of the putative corresponding *E* gene; the gene is likely to reside among the relatively smaller number of predicted targets in that genome that are shared by the two alleles.

Up to now 12 *R* genes against BLB have been cloned, of which 9 are triggered by TALEs (Jiang et al. 2020; Chen et al. 2021; Luo et al. 2021). This proportion not only highlights the crucial role of this effector family in the rice-*Xoo* pathosystem but also the relevance of using TALEs as probes to identify resistance sources and clone the underlying genes. This quest is facilitated by the ability to identify candidate targets of TALEs by combining EBE prediction and expression analysis, owing to the modular nature of TALE DNA interactions and their strong transcriptional upregulation of targets (Boch et al. 2014). In a proof-of-concept study pioneering this strategy for *E* gene identification, Strauss et al (2012) identified the pepper *Bs4C* gene, which is specifically induced by the TALE AvrBs4 from *X. axonopodis* pv. *vesicatoria*. More recently, a mix of map-based cloning and EBE predictions led to the isolation of the *E* gene *Xa7* in rice, which is triggered by the *Xoo* TALE AvrXa7 (Chen et al. 2021; Luo et al. 2021). In our study, out of the 86 accessions phenotyped, 12 were found to be resistant to both African *Xoo* strains MAI1 and BAI3, including 7 accessions of *O. sativa*, 2 of *O. glaberrima* and 3 NERICAs. Moreover, 12 *O. glaberrima*, 3 *O. sativa*, 3 *O. barthii* and 2 NERICA accessions were discovered to be resistant specifically to the Malian strain MAI1.

Among the 12 accessions identified as resistant to *Xoo* African strains MAI1 and BAI3, 9 accessions, including CT13432 and FKR47N, showed a phenotype typical of *Xa1* resistance (Table S2; Fig. S3). This is not surprising since *Xa1* and its functional homologs exhibit broad-spectrum resistance against African *Xoo* specifically, owing to the lack of iTALEs in examined African *Xoo* strains (Ji et al. 2016; Read et al.;2016). The *Xa1* allele that we discovered in the rice variety CT13432 appears to contain seven LRRs, like the allele originally identified as *Xa45(t)*, which was cloned from the wild rice variety *Oryza nivara-1* (Ji et al. 2020). Other alleles reported to date contain 4 (*Xa14*), 5 (*Xa2* and *Xo1*), and 6 repeats (*Xa1*). Variety FKR47N contains an allele that appears to have 6 LRRs. FKR47N, also called NERICA 17, is the result of crossing *O. glaberrima* variety CG14 and *O. sativa* spp. *japonica* variety WAB181-18, backcrossing to WAB181-18 as recurrent parent. *Xa1* alleles have been reported in other NERICAs. Alleles in NERICAs 5 and 7, the recurrent parent of which is the *O. sativa* spp. *japonica* rice variety WAB56-104, have five and six LRRs respectively. NERICAs 12 and 14, coming from the *O. sativa* spp. *japonica* rice variety WAB56-50, each carry an allele with five LRRs (Ji et al. 2020). According to Ji et al. (2020), a search for *Xa1* allelic members in the 3000 rice genomes revealed that approximately 15% contain the *Xa1* signature sequence. A phenotypic screening for *Xa1*-like resistance in 87 rice accessions revealed its in 16, including different rice species such as *O. glaberrima, O. nivara* and *O. sativa* (Ji et al. 2020). A complementary study on more than 500 *O. sativa* accessions revealed that *Xa1* alleles were present in *aus, indica*, temperate and tropical *japonica* (Zhang et al. 2020). The frequency of *Xa1* alleles among diverse rice varieties and the variability of LRRs number among *Xa1* alleles observed in these studies and revealed further by our results, underscores the unanswered question of the origin of the gene and the drivers of its diversification.

BLB represents a serious threat to rice production in Africa. Varietal resistance is the best strategy to control BLB durably, but it requires sources of resistance effective against the local pathogen genotypes. So far, no African strain of *Xoo* with iTALE/truncTALE has been identified, which makes *Xa1* and its alleles promising tools to control BLB in Africa. Although the durability of *Xa1* in Africa is difficult to predict, the risk of emergence of strains of *Xoo* originating from Asia and their spread through the African continent seems high in the current context of intense global exchanges of rice germplasm and insufficient phytosanitary measures. Pyramiding of resistance genes against BLB is of great value to extend the durability and the spectrum of resistance within a rice variety (Oliva et al. 2019). CT13432 combines several rice blast resistance genes (Pi1, Pi2 and Pi33) (Tharreau et al. 2007; Utami et al. 2011), but here we found out that it also carries at least two, distinct types of resistance to BLB, an apparent *Xa1* allele and a putative, TalI-dependent *E* gene, making it an excellent material for breeding to create improved varieties for Africa.

### Conclusions

In this study, we identified 12 rice accessions exhibiting resistance against African strains of *Xoo* and described 4 TAL effectors from these strains that exhibit avirulence activity. Analysis of the mechanisms underlying the resistance of the rice variety CT13432 revealed the occurrence of two overlapping sources of resistance including an apparent allele of the broad-spectrum resistance gene *Xa1* and an unidentified putative *E* gene that is activated by TalI. Combining transcriptomics and EBE prediction tools in the genome of CT13432 may allow discovery of this gene in the future. This approach would also be useful to decipher what genes in IR64 confer resistance against strains of *Xoo* with TalD.

## Material and Methods

### Bacterial Strains and Growth Conditions

Bacterial strains used in this study are listed in Table S4. *Escherichia coli* was cultivated at 37°C in liquid or solid (15 g agar per L) Luria-Bertani (LB) medium (10 g of tryptone, 5 g of yeast extract, and 5 g of NaCl per L of distilled water), and *Xanthomonas oryzae* pathovars at 28°C in liquid or solid (16 g agar per L) peptone sucrose medium (10 g of peptone, 10 g of sucrose, and 1 g of glutamic acid per L of distilled water).

### Plant Materials and Plant Inoculations

The rice accessions screened in this study are listed in Table S1. Plants used were grown in a greenhouse under cycles of 12h of light at 28°C with 80% relative humidity (RH) and 12h of dark at 25°C with 70% RH. When used for bacterial quantification assays, plants were grown in a growth chamber at 28°C and 80% RH (day and night). For lesion length assays, leaves of 5-week-old plants were inoculated by leaf-clipping with a bacterial suspension at an optical density at 600 nm (OD_600_) of 0.2. Lesion lengths were measured 15 days post inoculation (dpi) on at least eight leaves from individual plants per experiment. Leaves of 3-week-old plants were infiltrated with a needleless syringe and a bacterial suspension at an OD_600_ of 0.5. Symptoms were photographed at 5 dpi. For bacterial quantification assay, 5-week-old plants were inoculated by leaf-clipping and 10 cm distal leaf fragments of three leaves from three individual plants per condition were cut in half and ground in liquid nitrogen separately as reported previously (Yu et al. 2011).

### Plasmid Transformation

Plasmids were introduced into *E. coli* cells by heat-shock transformation and into *Xoo* by electroporation or triparental mating (Figurski and Helinski 1979). Appropriate antibiotics for selection were added to growth media at the following final concentrations: rifampicin, 100 μg.ml ^-1^; gentamicin, 100 μg.ml ^-1^; tetracycline, 100 μg.ml ^-1^; kanamycin, 100 μg.ml ^-1^. *tal* genes of the African *Xoo* strains MAI1 and BAI3 are cloned in pSKX1, which confers gentamicin resistance (Tran et al., 2018; Table S4). The truncTALE *tal2h* is cloned in pKEB31, which confers tetracycline resistance (Read et al. 2016; Table S4). The plasmids are compatible with each other.

### Mutant library construction and characterization

The BAI3*Δtal* mutants were generated upon electroporation of the wild-type strain BAI3 with the suicide plasmid pSM7 (Cernadas et al. 2014). Mutant strains were selected with kanamycin at 100 μg.ml ^-1^. The mutant library was characterized by Southern blot analysis. Extraction of *Xoo* genomic DNA (gDNA) was performed using the Wizard Genomic DNA Purification kit (Promega^®^). Four micrograms of gDNA were digested by *Bam*HI-HF (New England Biolabs) at 37°C overnight. The digested DNA was separated in a 1 % agarose gel at 50 Volts for 72 hours at 4°C and transferred to a nylon membrane (Roche^®^) overnight. Hybridization was conducted according to the procedures described in the DIG High Prime DNA Labeling and Detection Starter kit II protocol (Roche^®^), using a 725-bp C-terminal *TalC*_MAI1_ amplicon as probe (Yu et al. 2011).

### Construction of *talI* activation domain mutant

The *talI*_*MAI1*_ΔAD construct was made by swapping the central repeat region of a *talF*_*MAI1*_ΔAD construct with that of *talI*_MAI1_. To create the *talF*_*MAI1*_ΔAD construct, a PCR product corresponding to a 520 bp fragment encoding the C-terminus of *talF*_MAI1_ was obtained with primers ΔAD_Fw and ΔAD_Rv (Table S3), thereby allowing to introduce an inframe 167-nt deletion. This amplicon was introduced into pSKX1_*talF*_*MAI1*_ between the *Pvu*I and *Hind*III restriction sites using T4 DNA ligase (Promega^®^) according to manufacturer’s recommendations, creating pSKX1_*talF*_*MAI1*_ΔAD. The *talI*_*MAI1*_ central repeat region was then swapped into pSKX1_*talF*_*MAI1*_ΔAD using *Stu*I and *Aat*II (NEB, New England Biolabs) to generate pSKX1_*talI*_*MAI1*_ΔAD. The plasmid was confirmed by Sanger sequencing.

### Expression Analysis by Western Blotting

*Xoo* strains carrying the different combinations of TALE-encoding plasmid, truncTALE (*tal2h*) -encoding plasmid, and empty vectors (pSKX1 or pKEB31) were grown in liquid peptone sucrose medium supplemented with the corresponding antibiotics at 28°C. Cells of 1 ml of a bacterial suspension at an OD_600_ of 0.4 were harvested. Proteins were extracted with the BugBuster ^®^ Master Mix (Novagen), according to manufacturer’s recommendations. The total protein concentration of each sample was calculated by a Bradford assay following the Bio-Rad Protein Assay protocol (Bio-Rad, USA), and protein concentrations adjusted. TALE and truncTALE expression was analyzed by sodium dodecyl sulfate polyacrylamide gel electrophoresis using a 4-15% polyacrylamide gel and immunoblotting using the anti-TALE polyclonal antibody produced by Read et al. (2016) followed by a horseradish peroxidase conjugated rabbit secondary antibody (Sigma-Aldrich). Detection was carried out using the Thermo Scientific™ Pierce™ ECL 2 Western Blotting Substrate kit and a Typhoon™ FLA 9500 (General Electric Healthcare Life Sciences, USA) for imaging.

### PCR-amplification of putative functional *Xa1* alleles

To identify rice accessions harboring a potentially functional *Xa1* allele and to decipher the number of 93 aa repetitions, genomic DNA of rice accessions was extracted using an adaptation of the Murray and Thompson protocol (Murray and Thompson 1980) and was subjected to PCR using pairs of primers XaL-F1/XaL-R1 and XaL-F2/XaL-R2, respectively (Table S3; Ji et al. 2020).

## Author’s Contributions

ML, BS, and MH designed the experiments. ML and MH conducted the experiments. ET participated in plant material growth and propagation. ML, AB, BS, and MH performed data analysis and wrote the manuscript. All authors read and approved the final manuscript.

## Funding

This work was supported by the CRP-Rice (CGIAR Research Program) to MH and BS, and the Plant Genome Research Program of the National Science Foundation (Division of Integrative Organismal Systems IOS-1444511 to AJB).

## Availability of Data and Materials

The datasets supporting the conclusions of this article are provided within the article and its supplementary information files.

## Declarations

### Ethics Approval and Consent to Participate

Not applicable

### Consent for Publication

Not applicable

### Competing interests

The authors declare that no competing interests exist.

## Figure titles and legends

**Fig. S1. Molecular characterization of a library of BAI3Δ*tal* mutant strains**

Genomic DNA of the wild-type *Xoo* strain BAI3 and derivative BAI3*Δtal* mutants were digested by *Bam*HI*-*HF which cuts on either side of the central repeat region of *tal* genes and revealed by Southern blot using a 725-bp C-terminal *talC*_MAI1_ amplicon as probe (Yu et al. 2011). Individual mutants were obtained for each *tal* gene with the exception of *talH*. One double *talG/talH* mutant was analyzed in stead. BAI3Δ*talB* was obtained previously, also using the suicide plasmid pSM7 (Tran et al., 2018), and is therefore not included here. *tal* genes are indicated to the left and DNA sizes to the right. Red arrows indicate the *tal* gene(s) that were mutated.

**Fig. S2. Phenotypic responses upon leaf-infiltration of CT13432 and FKR47N plants with BAI3Δ*talI* and BAI3Δ*talF* mutants**.

Leaves of rice varieties IR64, CT13432, FKR47N and Azucena were infiltrated with the wild type African *Xoo* strain BAI3 and the mutant derivatives BAI3Δ*talD*, BAI3Δ*talI* and BAI3Δ*talF*. Inoculated leaves were photographed at 5 dpi.

**Fig. S3. The tal2h truncTALE reveals Xa1-like resistance against the Xoo strain BAI3 in several rice varieties**.

Leaves of rice accessions, including Carolina Gold Select which carries *Xo1*, and IRBB1 which carries *Xa1*, were infiltrated with *Xoc* strain BLS256 which naturally carries the *tal2h* truncTALE gene, the mutant strain BLS256Δ*tal2h*, as well as with *Xoo* strain BAI3 carrying an empty vector (EV) or *tal2h*. The Asian *Xoo* strain PXO99^A^ which harbors two truncTALEs, was used as an additional positive control. Leaves were photographed at 5 dpi.

**Fig. S4. Western-blot analysis of *Xoo* total protein extracts using an anti-TALE antibody**. Proteins extracts prepared from *Xoo* strain BAI3 carrying the empty vector pKEB31 (EV), BAI3Δ*talI* carrying the empty vector pKEB31 (EV), BAI3 carrying the *tal2h* truncTALE gene, and BAI3Δ*talI* with *tal2h* and the pSKX1 empty vector (EV), or pSKX1 containing *talI*_*BAI3*_ or *talI*_*MAI1*_ or *talI*_*MAI1*_ΔAD. BAI3 *tal* genes and *tal2h* are indicated to the left and right, respectively. Molecular weight is indicated.

## Notes

### Competing Interest Statement

The authors have declared no competing interest.

